# Identification of novel transcriptional regulators of PKA subunits in *Saccharomyces cerevisiae* by Quantitative Promoter–Reporter Screening

**DOI:** 10.1101/015891

**Authors:** C Pautasso, K Chatfield-Reed, G Chua, V Zaremberg, S Rossi

## Abstract

The cAMP dependent protein kinase (PKA) signaling is a broad specificity pathway that plays important roles in the transduction of environmental signals triggering precise physiological responses. cAMP-signal transduction specificity is achieved and controlled at several levels. The *Saccharomyces cereviciae* PKA holoenzyme consists of two catalytic subunits encoded by *TPK1*, *TPK2* and *TPK3* genes, and two regulatory subunits encoded by *BCY1* gene. In this work we studied the activity of these gene promoters using a reporter-synthetic genetic array screen, with the goal of identifying novel regulators of PKA subunits expression. Gene ontology (GO) analysis of the regulators identified showed that these regulators were enriched for annotations associated with roles in several GO biological process, as lipid and phosphate metabolism or transcription regulation and regulate all or some of the four promoters. Further characterization of the effect of these pathways on promoter activity and mRNA levels pointed to inositol, inositol polyphosphates, choline and phosphate as novel upstream signals that regulate transcription of PKA subunit genes. In addition, within each category there are genes that regulate only one of the promoters and genes that regulate more than one of them at the same time. These results support the role of transcription regulation of each PKA subunit in cAMP specificity signaling. Interestingly, many of the known targets of PKA phosphorylation are associated with the identified pathways, opening the possibility of a reciprocal regulation in which PKA would be coordinating different metabolic pathways and these processes would in turn, regulate expression of the kinase subunits.

## INTRODUCTION

The great variety of cellular processes regulated by the cAMP-protein kinase A (PKA)-pathway must be strictly controlled to maintain specificity in the response. Different regulatory mechanisms must exist to ensure the phosphorylation of the correct substrate in response to the proper stimulus. Several combined factors determine the differential effects of the signaling cascade initiated by cAMP. In mammals, different biochemical properties and substrate specificity displayed by PKA isoenzymes and their localization through the association with anchoring proteins (AKAPs) have been shown to contribute to specificity in the cAMP pathway (Skålhegg and Taskén 1997). Transcriptional regulation of the PKA subunits and their level of expression should also have a critical impact on signal specificity, but little is known about how these processes are regulated. The regulatory (R) and catalytic (C) subunits of mammalian PKAs are transcriptionally regulated by hormones and mitogens acting through different receptors like G-protein coupled receptors (Jahnsensg *et al.* 1985; Pariset *et al.* 1989; Landmark *et al.* 1993a, 1993b) or tyrosine kinases-associated receptors (Skålhegg and Taskén 1997). PKA subunits have shown differential expression patterns at different developmental and differentiation stages as well as in different tissues (Cadd and Mcknight 1989; Beebe *et al.* 1990; Hougel 1992; Landmark *et al.* 1993; Reinton *et al.* 1998; Cumming *et al.* 2007). cAMP positively modulates transcription of PKA subunits involving stabilization of their mRNAs as well as on both R and C protein stability after dissociation of the holoenzyme (Houge *et al.* 1990; Taskén *et al.* 1991; Knutsen *et al.* 1991; Hougel 1992).

In the model organism *Saccharomyces cerevisiae*, PKA controls a variety of essential cellular processes associated with fermentative growth, entrance into stationary phase, stress responses and development (Thevelein and Winde 1999; Thevelein *et al.* 2008; Smets *et al.* 2010). This pleiotropic role of PKA also needs a tight regulation. The structure of the PKA holoenzyme is conserved from mammals to yeast, and consists of a heterotetramer composed of a regulatory subunit homodimer and two associated catalytic subunits. The yeast catalytic subunits are encoded by three genes, *TPK1*, *TPK2* and *TPK3*, and the regulatory subunit, by the *BCY1* gene. Among the factors contributing to the cAMP-PKA pathway specificity in yeast are the synthesis, breakdown and spatial localization of cAMP, Tpk isoform-dependent phosphorylation of substrates and subcellular localization of the holoenzyme (Griffioen and Thevelein 2002; Vandamme *et al.* 2012; Engelberg *et al.* 2014). We are interested in understanding how transcriptional regulation of the PKA subunits contributes to the specificity of the cAMP-PKA signaling. We have previously conducted investigations aimed at characterizing the promoter activity of the *BCY1* and *TPK* genes and have demonstrated that the promoter of each isoform of *TPK* and of *BCY1* is differentially regulated during growth phase and stress conditions (Pautasso and Rossi 2014). *TPK1* promoter activity is positively regulated during heat shock and saline stress but *TPK2, TPK3*, and *BCY1* promoters, unlike *TPK1*, are not activated under these stress conditions. Therefore the expression of each PKA subunit involves different mechanisms in response to heat shock or saline stress. However, the four promoters of PKA subunits share an inhibitory autoregulatory mechanism since all of them are downregulated by PKA activity (Pautasso and Rossi 2014). Taking into account these antecedents, our aim in this work was to identify novel transcriptional regulators of PKA subunits. Taking advantage of the unique tractability of yeast, we used an unbiased high-throughput approach to uncover regulators of the promoters of the *BCY1* and *TPK* genes. We performed a reporter-synthetic genetic array (R-SGA) screen previously described to assess the effect of viable deletion mutants on the transcription of *TPK1, TPK2, TPK3* and *BCY1* promoters. The R-SGA screen makes use of a two-colour promoter-reporter system that is delivered to the array of viable haploid deletion mutants using high-throughput genetics (Kainth *et al.* 2009). This reporter-based screen has shown to be a powerful strategy for identification of regulatory proteins and upstream signals involved in promoter regulation (Kainth and Andrews 2010). Using this approach, we were able to identify unique pathways that differentially regulate the activity of the *BCY1, TPK1, TPK2* and *TPK3* promoters. Clustering analysis of the genes identified revealed enrichment in genes with roles in several GO biological process. From these GO, lipid and phosphate metabolism, and regulation of transcription categories were further characterized and validated using β-galactosidase reporter assays and qRT-PCR. The results of our genetic screen pointed to inositol, choline and phosphate as novel upstream signals that regulate transcription of PKA subunit genes.

## EXPERIMENTAL PROCEDURES

### Strains, plasmids and culture conditions

Table 1 lists the genotype of the strains used in this study. For the β-galactosidase reporter assays and RNA purification the strains were cultivated at 30° to early log phase (OD_600_=1) or late log phase (OD_600_=3.5) in synthetic SD media containing 0.67% yeast nitrogen base without amino acids, 2% glucose plus the necessary additions to fulfill auxotrophic requirements. In the assays to measure inositol and choline effects, yeast cultures were grown at 30° in a complete synthetic medium lacking inositol, choline (Klig and Henry 1984) and uracil (in the case of reporter plasmids). Where indicated, 75 µM inositol (I) and/or 1 mM choline (C) was added. The assays to analyze the phosphate effect were performed in low-Pi medium containing 0.15 mM KH_2_PO4, and high-Pi medium containing 7.35 mM KH_2_PO_4_. High-throughput functional assays were performed with plasmids containing the 5’ regulatory region and nucleotides of the coding region of *TPK1, TPK2, TPK3* and *BCY1* genes (positions −800 to +10 with respect to the ATG initiation codon in each case), cloned into pBA1926 controlling GFP expression. The plasmids used in β-galactosidase reporter assays to measure the promoter activities were derived from the YEp357 plasmid (Myers *et al* 1986). The *TPK1-lacZ*, *TPK2-lacZ*, *TPK3-lacZ*, and *BCY1-lacZ* fusion genes contain the same 810 bp regulatory fragments included in the pBA1926 constructs.

**Table 1.**
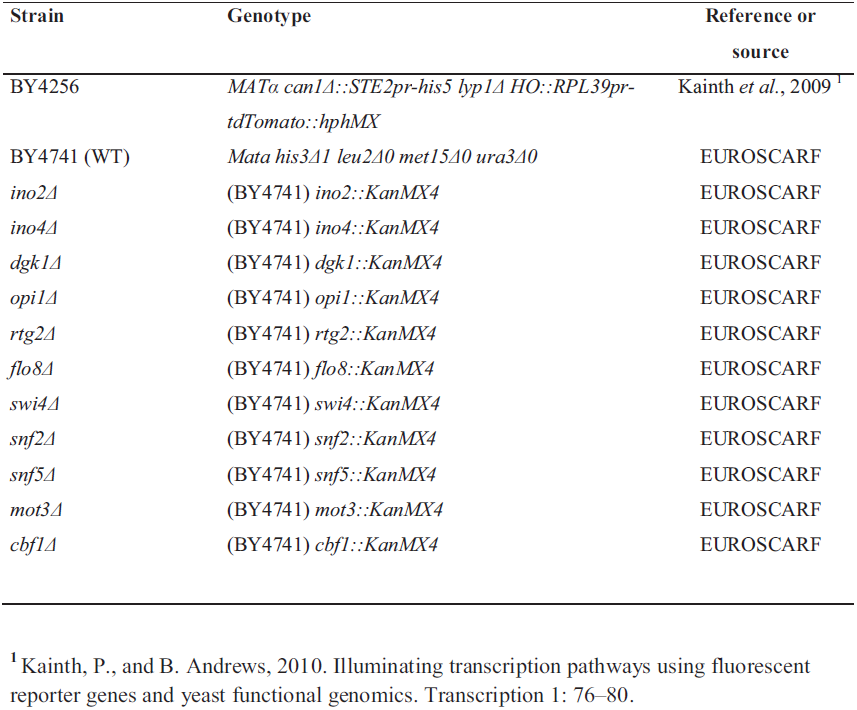
YEAST STRAINS

### High-throughput functional assay

The *Saccharomyces cerevisiae* array of 4500 viable strains, each missing a non-essential gene (BY4741 background, Euroscarf) was used in the R-SGA screen (Kainth *et al* 2009) to detect genes that affect the expression of a GFP reporter system under the control of *TPK1, TPK2, TPK3* and *BCY1* promoters The −800 to +10 sequences from each promoter were cloned upstream the GFP reporter gene. The constructs were used to transform the BY4256 strain carrying the RPL39pr–tdTomato reporter (RFP). After mating the collection strain with the BY4256 MATα strain carrying each of the *pr*GFP constructs, diploids were selected and sporulated followed by the selection of MATa xxxΔ haploids containing the *pr*GFP plasmid. Robotic manipulation of the collection was performed using a Biomatrix robot and plate imaging system (S&P Robotics Inc.). Colony size for each arrayed mutant was analyzed and positions on the array with no or slow colony growth were eliminated from further analysis. Colony fluorescence was assayed following 1 and 4 days of incubation on minimal glucose medium using PharosFX Molecular Imager (Bio-Rad). The data was analyzed with Array Gauge V1.2 software. The log2 GFP/RFP ratio from each colony on the array was calculated as described (Kainth *et al.* 2009). A large number of deletion mutants were represented twice in the array, and each screen was performed in duplicate, resulting in quadruplicate measurements for some deletion mutants. These log2 ratios were transformed to robust Z scores using median and median absolute deviation, and p-values were assigned to these Z scores based on a normal distribution. A p-value cut-off of 0.01 and 0.05 was set. A list of 260 deletion mutants that usually appear as hits in these screens (Henny Goettert and Brenda Andrews, personal communication), was removed from further analysis of specific *TPKs* and *BCY1* promoter regulators.

### GO analysis

The distribution of regulators into functional groups was assessed by measuring the enrichment for genes in the same functional category according to the gene annotation database of FunSpec (http://funspec.med.utoronto.ca/). The enrichment of the genes belonging to the GO Biological Process, GO Molecular Function and MIPS functional classification categories was calculated respect to the total gene number and the total number of genes belonging to the GO category, with a p-value of 0.05 or 0.01. The hypergeometric distribution was used to calculate a p-value for this fraction, and took p-value <0.05 or p<0.01 to be significant. Only the results for stationary growth phase are shown.

### β-galactosidase assays

Cells were grown on SD medium at 30° up to an OD_600_ of 3.5. Aliquots (10 ml) of each culture were collected by centrifugation and resuspended in 1 ml buffer Z (60 mM Na_2_HPO_4_, 40 mM NaH_2_PO_4_, 10 mM KCl, 1 mM MgSO_4_). β-galactosidase activity was measured and expressed as Miller units (Miller 1972). Results shown correspond to a representative experiment which was repeated at least three times, each one performed in duplicates or triplicates.

### qRT-PCR

Total RNA was prepared from different yeast strains, grown at 30° to the same OD_600_ as for β-galactosidase assays, using standard procedures. To determine the relative levels of specific *TPK1, TPK2, TPK3* and *BCY1* mRNAs, a quantitative RT-PCR experiment was carried out. Aliquots (∼10 µg) of RNA were reverse-transcribed into single-stranded complementary cDNA using an oligo-dT primer and Superscript II reverse transcriptase (Life Technologies). The single-stranded cDNA products were amplified by real time PCR using gene-specific sense and antisense primers (mRNA *TPK1:* Fw: 5’ CCGAAGCAGCCACATGTCAC 3’, Rv: 5’ GTACTAACGACCTCGGGTGC 3’; mRNA *TPK2:* Fw: 5’ GCTTGTGGAGCATCCGTTTC 3’, Rv: 5’ CACTAAACCATGGG TGAGC 3’; mRNA *TPK3:* Fw: 5’ CGTTGGACAAGACATTCCTG 3’, Rv: 5’ GTCGGT TATCTTGATATGGCC 3’; mRNA *BCY1:* Fw: 5’ CGAACAGGACACTCACCAGC 3’, Rv: 5’ GGTATCCAGTGCATCGGCAAG 3’; mRNA *TUB1*((α-Tubulin gene): Fw: 5’ CAAGGGTTCTTGT TTACCCATTC 3’, Rv: 5’ GGATAAGACTGGAGAATATGAAAC 3’). The PCR products were visualized using SYBR® Green (Life Technologies). The relative mRNA levels of *TPK1, TPK2, TPK3* and *BCY1* were first normalized to those of *TUB1*and then compared to each other. Quantitative data were obtained from three independent experiments and averaged.

## RESULTS and DISCUSSION

### Validation of the R-SGA genetic screen

We used the Dual-Reporter Functional Genomic Screen, R-SGA approach (Kainth *et al.* 2009) to survey the viable deletion collection for mutants that affect the transcription of promoters of the PKA subunits, (*prTPK1*-GFP, *prTPK2*-GFP, *prTPK3*-GFP and *prBCY1*-GFP). Differential GFP expression was assessed by scanning fluorescence intensities (GFP and control RFP) directly from colonies arrayed on agar plates obtaining in this way a genome-wide result of the effect of viable deletion mutants on the activity of the promoters of PKA subunits. The R-SGA assay was performed from colonies arrayed in defined media grown for four days with glucose as carbon source. The GFP:RFP ratios were normalized and the log2 values calculated as described (Figure S1). Decreased GFP:RFP ratios correspond to deletion of activators while increased ratios reveal deletion of putative repressors (Kainth *et al.* 2009). We considered the possibility that the genome-wide screen gave false positives using a p<0.05, however our control was the cAMP-signaling pathway as regulator. Although not much is known about transcriptional regulation of these genes, we have previously demonstrated that *TPK*s promoters are inhibited by PKA activity, and that Tpk2 had a stronger inhibitory effect on *TPK1* and *TPK3* promoters when cells were grown in liquid cultures (Pautasso and Rossi, 2014). Thus, we expected the deletion of *TPK2* to result in increased *prTPK1*-GFP and *prTPK3*-GFP expression compared to its effect on the control RPL39pr-RFP gene. We found that *TPK2* deletion did in fact cause a defect in *TPK1* transcription (Table S1). Reducing the cutoff to 1%, Tpk2 was not detected; thus, even though the false positives perhaps could be reduced using a lower cut off, this lower p-value could discard true regulators. The putative transcriptional regulators identified when the cut off was 0.05 are summarized in Table S1.

The R-SGA analysis was also performed from colonies arrayed on defined media with glucose as carbon source grown only one night instead of four days (data not shown). The results in this case showed a higher number of negative regulators and a lower number of positive regulators than those identified from plates grown four days (Figure 1). This is in agreement with previous results from our laboratory indicating that all the promoter subunits are upregulated during stationary growth phase (Pautasso and Rossi 2014). Here, although strains were grown in solid media plates instead of liquid cultures, the results showed the same tendency when comparing plates incubated for different periods of time.

**Figure 1.**
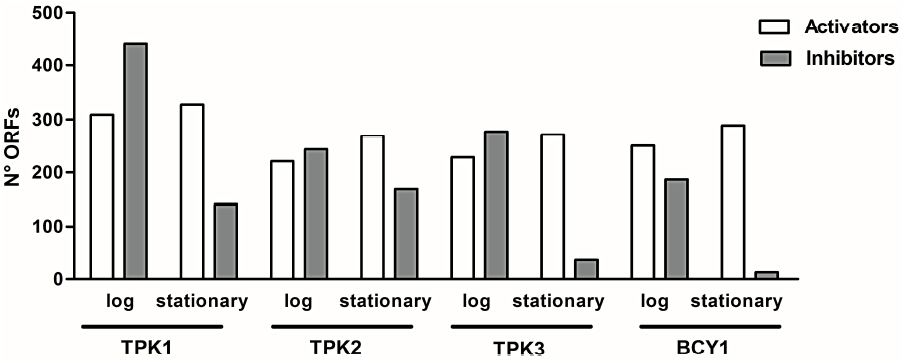
Regulators, activators or inhibitors, identified in the high-throughput functional assay from plates grown 1 (early log phase) or 4 (late log phase) days of incubation.

Altogether, the results of our screen were consistent with previous findings, and successful identifying some of the expected modulators, therefore validating the use of this approach for the unbiased identification of novel regulators of the expression of PKA subunits.

### Novel transcriptional regulators of PKA subunits

We focused our analysis on the results obtained from R-SGA screens performed from colonies arrayed on defined medium with glucose as carbon source grown four days at 30°. The log2 ratios were transformed to Z score and p-values assigned on the basis of a normal distribution (see Materials and Methods). This analysis identified 469 putative transcriptional regulators for *TPK1*, 437 for *TPK2*, 307 for *TPK3* and 299 for *BCY1* (Table S1).

Clustering analysis of the genes identified revealed a discrete number of gene ontology (GO) categories significantly enriched in each screen. The categories that were found to be enriched for *TPK1, TPK2*, *TPK3* and *BCY1* promoters with a cutoff of a p-value < 0.05 are listed in Figure 2 and Table S1. The results showed that several GO categories differentially affected the expression of *TPKs* and *BCY1*, while many others were common to all (Table 2).

**Figure 2.**
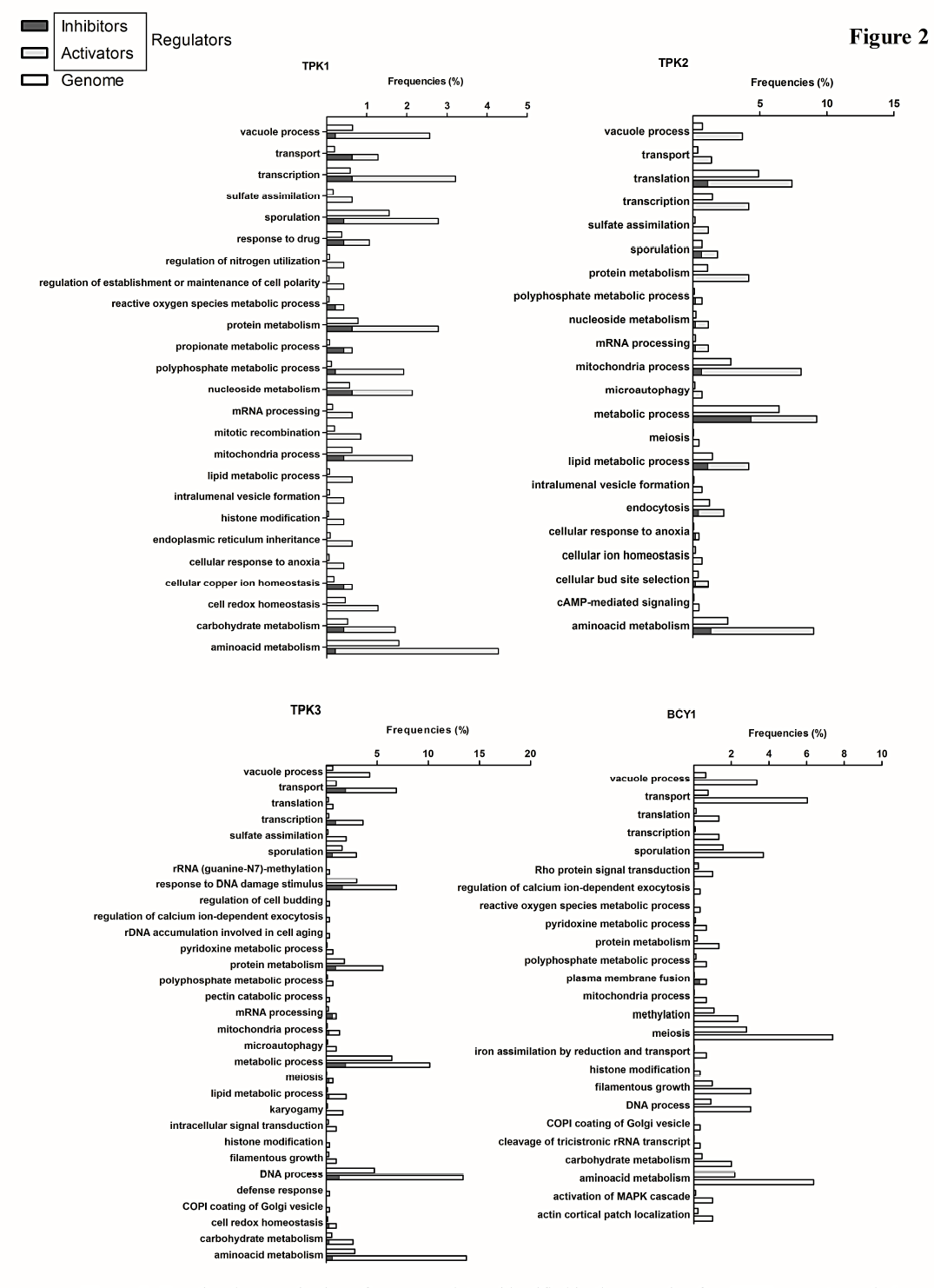
Functional categorization of genes regulators identified in the screening for *TPK1* promoter and of all genes in *S.cerevisiae.* Classifications were performed based on same functional category according to the gene annotation database of FunSpec (http://funspec.med.utoronto.ca/). Black and white bars represent the genome gene fraction in this category and the regulators fraction identified respectively.

**Table 2.**
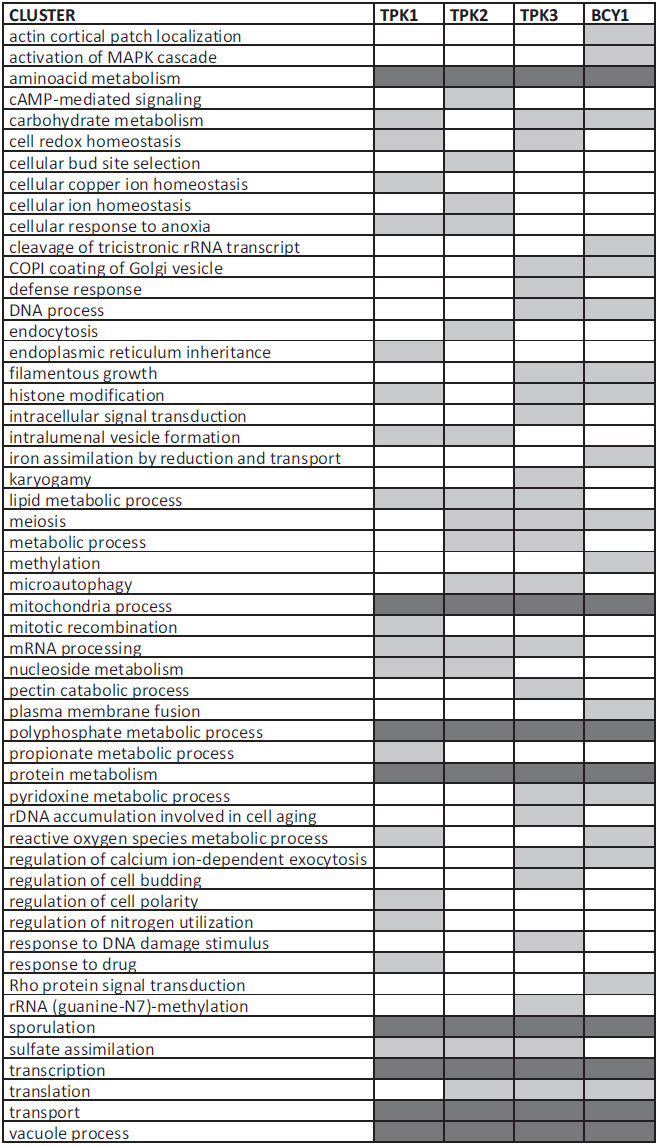
CLUSTERING

Using a more stringent cutoff of 0.01 the list of categories regulating PKA promoter activities was reduced from 42 to 9 (*TPK1*, 21,4%) 43 to 18 (*TPK2*, 41,8%) 86 to 19 (*TPK3*, 22%) and 57 to 7 (*BCY1*, 12,2%) (Table S1).

Overall, the results of the R-SGA screens have unveiled novel regulators of the PKA subunits promoter activities. Different biological process affect different each PKA subunit (twenty six categories), although some of them are shared for two (twelve categories), three (eight categories) or all the subunits (seven categories) (Figure S1). Even more making a detail analysis of all the genes included in the GO categories that were identified affecting all the subunits or at least the catalytic subunits we could observe that not all the same genes included in the GO category are identified as regulators. As is shown in the Figure 3, the Venn diagrams clearly show this result. Taking as a whole these results strengthen our hypothesis about the role of transcription regulation to collaborate with cAMP-PKA specificity signaling.

**Figure 3.**
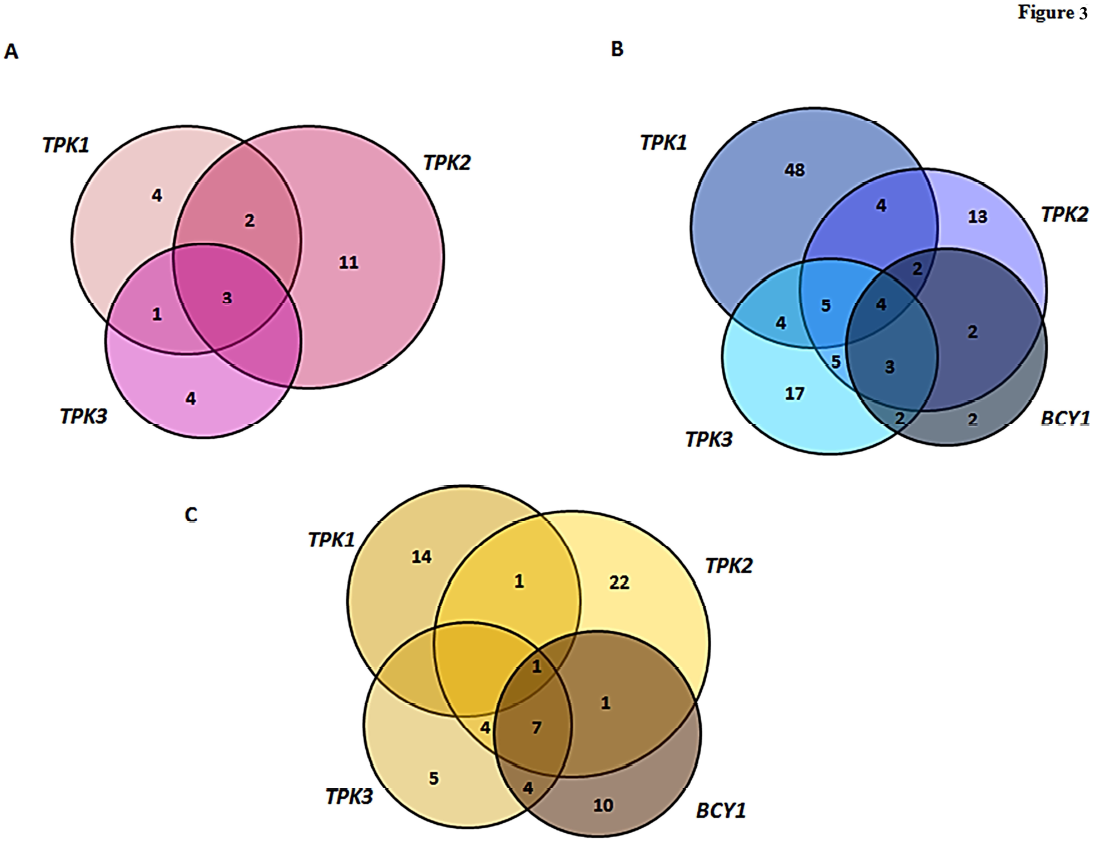
Venn diagrams including the regulators genes identified and indicating the common and differential regulators for each PKA subunit. Each diagram corresponds to one of the GO process analyzed: A) Lipids metabolism, B) Phosphate metabolism and C) Transcription regulation.

Interestingly two interrelated categories, lipid metabolism and phosphate metabolism were differentially enriched in the four screens (Figure 2). Lipid metabolism appeared as modulator of only *TPKs* promoter activities (not *BCY1* promoter), while Pi metabolism as regulator of *TPK2*, *TPK3* and *BCY1* promoters (Figure 3).

Lipids are major constituents of structural assembly of the cell and are integral to the growth, development and maintenance of the organism. In *S.cerevisiae*, lipid metabolism is altered by growth phase, by supplementation with inositol or choline and this regulation is controlled by genetic and biochemical mechanisms (Carman and Henry 2007). The regulation of genes related to phospholipid biosynthesis in *S. cerevisiae* through cis-acting upstream activating sequence (UAS_INO_) by inositol supplementation is thoroughly known. In addition, a role for inositol-polyphosphates, product of the reaction catalysed by the phosphatidylinositol-specific phospholipase C (Plc1) in the direct regulation of lipid biosynthesis has been proposed (Rupwate *et al.* 2012).

Lipids were only considered for years as building blocks for biological membranes, without any other specific functions. However, this idea has changed during the last two decades since certain classes of lipids are demonstrated to be involved in intracellular signaling processes or work as sensors of the membrane status of a cell. Among the lipids with important functions, phosphatidylinositol (PI) and derivatives are known to be involved in transduction of intracellular signals (Lemmon 2003) besides their role in vesicle trafficking (Odorizzi *et al.* 2000). Degradation products of phospholipid hydrolysis, phosphatidic acid (PA), diacylglycerol (DAG) and fatty acids, as well as sphingolipids, have also been shown to serve as messenger molecules. The intracellular amount of PA triggers the transcriptional regulation of genes containing the UAS_ino_. These genes are regulated by the transcription factors Ino2, Ino4, and the repressor Opi1. The transcriptional regulation of UAS_ino_ genes is generally controlled by the rate of phospholipid synthesis. (Henry *et al.* 2012). Under certain growth conditions (exponential phase, inositol depletion), the levels of PA are relatively high (by the action of several enzymes, among them Dgk1), and the Opi1 repressor is tethered to the nuclear/ER membrane through interaction with the integral membrane protein Scs2, an association is stabilized by interaction with PA. In this condition, UAS_INO_-containing genes are maximally expressed by the Ino2-Ino4 activator complex. In stationary phase, or with inositol supplementation, the levels of PA are reduced, Opi1 dissociates from the nuclear/ER membrane, and enters into the nucleus where it binds to Ino2 and attenuates transcriptional activation by the Ino2– Ino4 complex.

Among the genes identified in the GO category of lipid metabolism (summarized in Table S1), were many of the genes mentioned previously. these included genes known to be regulated by inositol and choline, such as *INO1*, *CHO2*, *SCS2*, *EKI1, FEN1*,and the transcription factors *INO2* and *INO4*, and genes involved in inositol polyphosphate metabolism including *PLC1* and those encoding the inositol kinases *ARG82, IPK1* and *KCS1.* The enrichment of this category survived was also detected with a more stringent cutoff of p<0.01,. In addition, genes coding for the phosphoinositide phosphatases *SAC1* and *INP51* as well as genes involved in sphingolipid metabolism like *ISC1* coding for the inositol phosphosphingolipid phospholipase C and *ELO2* and *ELO3* coding for fatty acid elongases involved in sphingolipid synthesis were identified. It is worth noting that phosphatidiylinositol (PI) metabolism in yeast is directly linked to the synthesis of sphingolipids as PI donates the phosphoinositol head group that is combined with ceramide for the production of complex sphingolipids.

Altogether these results pointed to a possible novel role of inositol as an upstream regulator of PKA subunits expression. To test this *TPK1*, *TPK2*, and *TPK3* promoter activities were assessed using promoter-lacZ-based reporter assays, in wild type cells (strain BY4741) grown to late log phase (OD_600_ 3.5) in minimal medium containing glucose but lacking or supplemented with 75 µM inositol and 1mM- choline (Figure 4A, upper panel). In fact, the activity of all three promoters were regulated by the presence of inositol. Interestingly, addition of choline to inositol depleted medium also activated these promoters and no major differences were found when inositol and choline were added together. This is somehow surprising as the effect of exogenous choline is usually much less dramatic than the effects of exogenous inositol in UASino containing genes (Carman and Henry 2007). In addition, no classic UASino sequences can be detected in any of the promoters analyzed. Addition of choline is known to significantly alter phosphatidylcholine turnover and the generation of the signalling lipids phosphatidic acid and diacylglycerol (Dowd *et al.* 2001)Altogether this may suggest that the effect of inositol and choline on the *TPKs* promoter activities is indirect by modulating the levels of signaling lipids and signaling soluble products of phospholipid turnover like inositol polyphosphates. (Lee *et al.* 2008)

**Figure 4.**
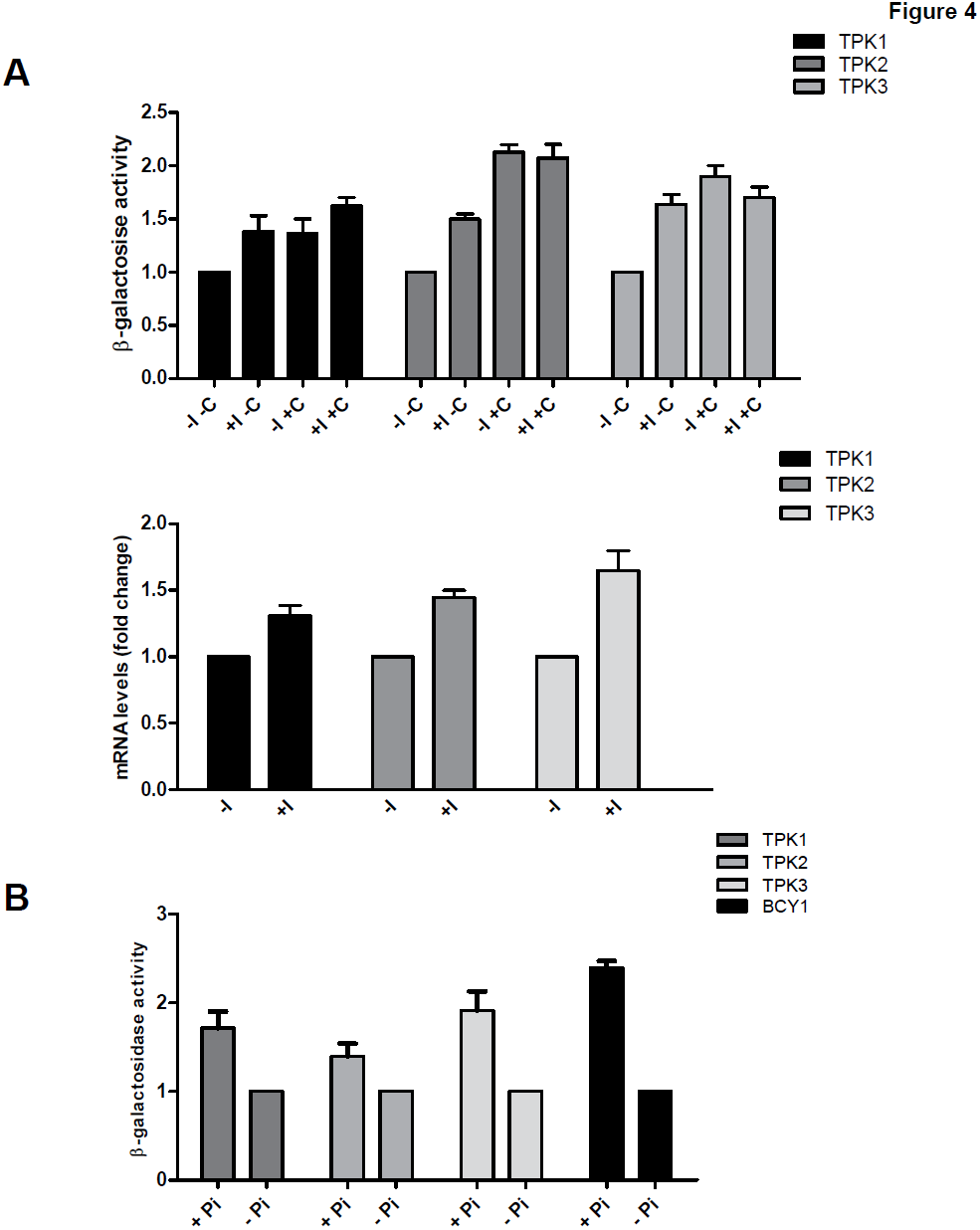
**A)** Upper panel, β-galactosidase activity was determined in WT cells (strain BY4741) carrying the *TPKs*-lacZ or *BCY1*-lacZ fusion gene. Cells were grown up to OD_600_=3.5 in minimal medium glucose lacking or supplemented with I and C. β-galactosidase activity is expressed in Miller Units. Results are expressed as the mean ± SD from triplicates within a representative assay and normalized to the *TPK1* values. Lower panel, *TPKs* and *BCY1* endogenous mRNA levels were determined in WT strain (BY4741) grown in the same conditions described above. The values were normalized to *TUB1* mRNA. The mRNA level for each subunit in the WT strain was defined as 1. **B)** β-galactosidase activity was measured as in A but using minimal medium with high phosphate (Pi) or low Pi.

PKA subunit promoters were activated and not repressed by inositol. In fact, many genes involved in phospholipid metabolism in yeast are repressed in response to inositol. For example, INM1 gene, encoding inositol 3-phosphate phosphatase, DPP1 and PAH1 genes, encoding lipid phosphate phosphatases and PIS1 gene, encoding phosphatidylinositol synthase are derepressed in the presence of inositol and in stationary phase ((Murray and Greenberg 1997; Oshiro *et al.* 2000; Jani and Lopes 2008). In addition to these, microarray analysis has also revealed many additional genes, that are regulated in response to the presence or absece of inositol and choline which are not involved in lipid metabolism (Santiago and Mamoun 2003; Jesch *et al.* 2005). In order to understand the mechanism involved in the regulation of PKA subunits expression we further tested the effect that deletion of various genes forming the inositol/PA sensing core had on the promoter activities of the PKA subunits. Reporter assays using β-galactosidase activity for each PKA subunit promoters were measured in mutant strains Δ*ino2*, Δ*ino4*, Δ*scs2*, Δ*opi1* and Δ*dgk1* (Figure 5A). The results indicated that the lack of *INO2, INO4* or *DGK1* upregulated *TPKs* promoters. On the contrary, in the Δ*opi1* strain *TPK1, TPK2* and *TPK3* promoters were downregulated. Taking into account these results we suggest that the regulation by inositol on *TPKs* promoters is indirect, and that Ino2/4 may regulate another gene that has repressor activity on *TPKs* promoters. To validate the results with another approach, the same mutant strains lacking the genes encoding regulatory proteins of the phospholipid biosynthetic pathway were used to measure the endogenous mRNA levels of each PKA subunit by RT-qPCR. The results showed that the mRNA levels of each subunit were not completely in agreement with promoter activities (Figure 5B), In the the Δ*no4* strain for *TPK1* and *TPK2* promoters, and Δ*ino2* strain for the three *TPKs* promoters, the mRNA levels were opposed to the promoter activity. Converse trends were also found for Δ*opi1.*

**Figure 5.**
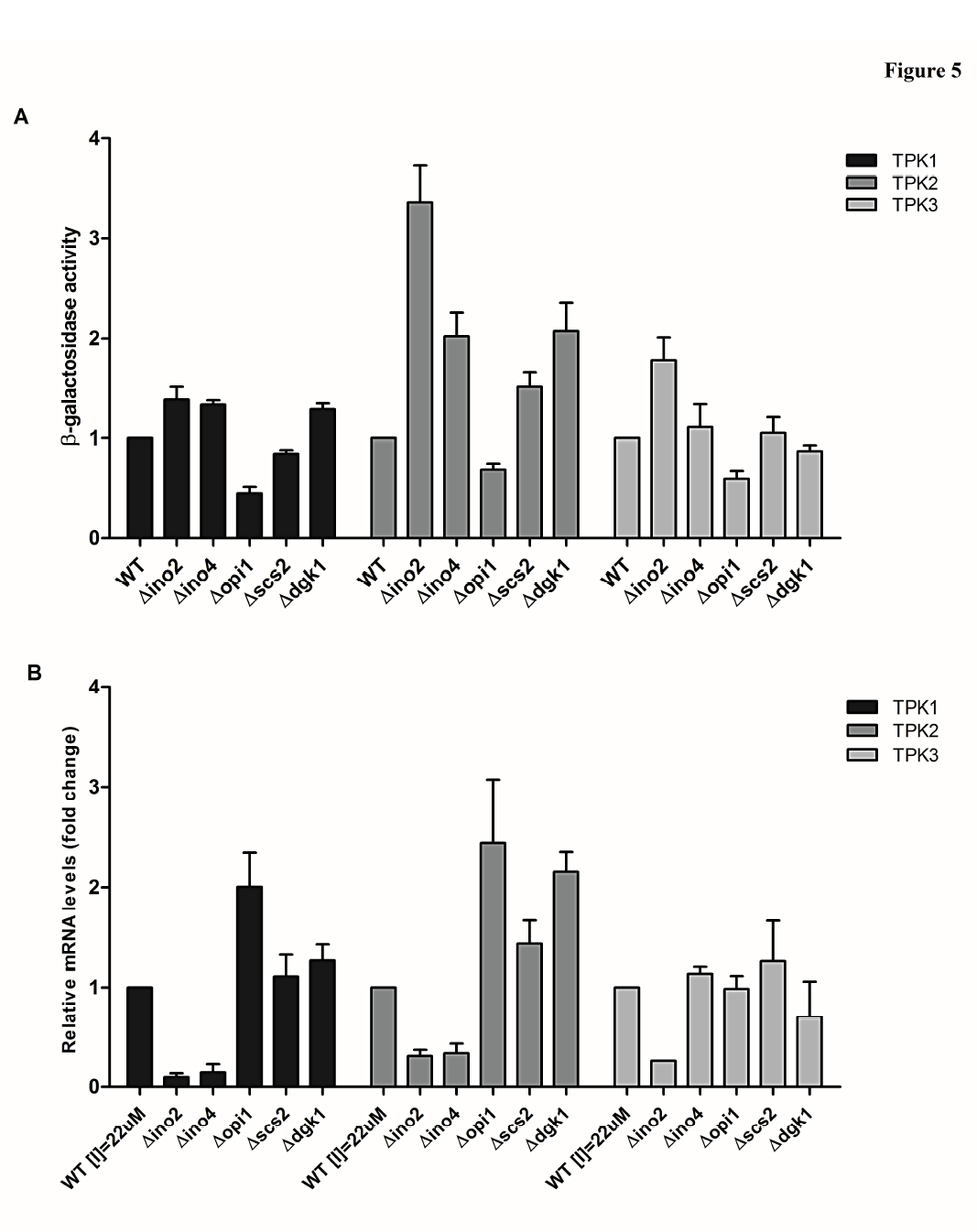
**A)** β-galactosidase activity was determined in WT (BY4741) *opi1*Δ, *ino2*Δ, *ino4*Δ, *scs2*Δ or *dgk1*Δ strains carrying the *TPKs*-lacZ or *BCY1*-lacZ fusion gene. The results are expressed in Miller Units and as the mean ± SD for replicate samples (n=4) from independent experiments. **B)** *TPKs* and *BCY1* endogenous mRNA levels were determined in WT (BY4741), *opi1*Δ, *ino2*Δ, *ino4*Δ, *scs2*Δ or *dgk1*Δ strains by qRT-PCR and normalized to *TUB1* mRNA. The mRNA level for each subunit in the WT strain was defined as 1.

These results, although unexpected, were not surprising since it was reported some genes involved in lipid metabolism that are regulated at the level of mRNA stability *INO4, INO1* and *CHO2* genes, which are also involved in the phospholipid biosynthetic pathway, are regulated at the level of mRNA stability by inositol and choline (Robinson and Lopes 2000). The inverse relation with reporter assays in Opi1 mutant strain suggest that Opi1 could regulate the PKA subunits expression at the transcription level and mRNA stability as it was already proposed for the regulation of *INO4* expression (Robinson and Lopes 2000).

This type of regulation has been further supported by the discover of synthegradase factors that modulate not only the transcription but also the decay of mRNA as a two-arm mechanism to be more responsive to regulatory signals (Henry *et al.* 2012). Recent studies have demonstrated that environmentally induced genes are subject to transcriptional upregulation along with an increase in decay rate of the transcripts (Bregman *et al.* 2011). This coupling almost always involves enhancement of both mRNA synthesis and decay or, conversely, repression of both mRNA synthesis and decay (Shalem *et al.*; Molin *et al.* 2009; Elkon *et al.* 2010; Rabani *et al.* 2011). Finally it has been proposed that an important proportion of the yeast genes examined is likely to be regulated by synthegradases during optimal proliferation conditions and the number of genes is likely to increase in stress conditions (Dori-Bachash *et al.* 2011). All these antecedents would indicate an important role of promoters, transcription factors and synthegradases in the regulation of both synthesis and decay of the PKA subunits mRNAs.

Another GO category which showed significant gene enrichment was phosphate metabolism including PHO80, PHO81, PHO84, PHO85 and PHO87 among other genes. The complex formed by the regulatory proteins Pho80, Pho81 and the cyclin kinase Pho85 are known to regulate the response to phosphate limitation (Huang et al. 2007) Similar to inositol and choline the presence of high phosphate (7.35 mM KH2PO4) increased the activity of all promoters compared to low phosphate (0.15 mM KH2PO4) (Figure 4B). Therefore promoter activity was lower in conditions of phosphate limitation. The effect of phosphate may be linked to its regulatory role on inositol metabolism and the generation of inositol polyphosphates, which are known regulators of gene expression in the phosphate sensing pathway (Lee *et al.* 2008).In addition, it was reported that phospholipid biosynthesis is coordinated with phosphate utilization via the bHLH proteins, Ino2/4 and Pho2/4 (He *et al.* 2012). We have observed that the PKA subunit promoters responded to the presence of inositol, choline and also Pi, suggesting a possible cross-regulation between both pathways.

The repression of genes containing UAS_INO_ is considered to be a mechanism for preferentially synthesizing large amounts of PI by inhibiting phosphatidylserine (PS) synthase activity, thereby making CDP-DAG available for PI synthase (Kelleys *et al.* 1988). This in turn, would lead to PI phosphorylation and formation of phosphoinositides Breakdown of PI(4,5)P_2_ gives rise to a pool of inositol-polyphosphates with important roles in the control of gene expression (Rupwate (Banfic *et al.* 2013; Galdieri *et al.* 2013). We have found that several genes included in the pathway of biosynthesis of inositol-polyphosphates were functionally enriched (p<0.01), indicating the possible importance of these phosphoinositides in the regulation of the activity of PKA subunit promoters. The hydrolysis of PI(4,5)P_2_ by Plc1 yields two prominent eukaryotic second messengers: 1,2-diacylglycerol (DAG) and inositol 1,4,5-trisphosphate (IP3) (Divecha and Irvine 1995; Carman and Han 2011). Plc1 and four inositol polyphosphate kinases (Ipk2p/Arg82p, Ipk1p, Kcs1p, and Vip1p) constitute a nuclear signaling pathway that affects transcriptional control (Odom *et al.* 2000), export of mRNA from the nucleus (Monserrate and York 2010), homologous DNA recombination, cell death, and telomere length (Banfic *et al.* 2013). All of these genes have been identified in our screen as regulators of PKA subunit promoters. We are further investigating the role of this important lipid signaling pathway in the regulation of PKA subunits expression.

The synthesis of phospholipids is regulated by controlling the expression of enzymes and/or by modulating its catalytic activities. The protein kinases known to regulate the function of catalytic and regulatory proteins in the phospholipid synthesis include protein kinase A, protein kinase C, casein kinase II, and cyclin-dependent kinase. Phospholipid synthesis enzymes, which are regulated by phosphorylation, include PS synthase, CTP synthetase, choline kinase, and PA phosphatase. The transcriptional repressor Opi1 is also regulated by phosphorylation by PKA. Is interesting to notice that our results indicate the possibility of a regulatory loop in which the expression of PKA subunits is controlled by lipid metabolism pathway and PKA activity regulates in turn, enzymes involved in phospholipids synthesis by phosphorylation. The same possible effect was proposed above in the case of a reciprocal regulation among PKA coordinating respiratory function and the respiratory metabolism upregulating PKA subunit promoter activities. And in the same line of thought is the autoregulatory loop by which PKA activity regulates the promoter activities of the enzyme subunits.

Finally we analyzed a group of transcriptional regulators that arose from the screen, including transcription factors as Swi4, Cbf1, Rtg3, Mot3 and Flo8, and chromatin remodelers Snf2 and Snf5, employing the reporter gene methodology in mutant strains (Figure 6). The results demonstrate the different effect that the absence of each transcription regulator caused on each PKA subunit promoter. The case of Flo8 transcription factor is noteworthy since it was identified as regulator of *TPK3* and *BCY1* transcription in our R-SGA screens even though the colonies in the array have a BY4741 background. It is well known that yeast strains derived from S288c, like BY4741, are incompetent for filamentous growth due to a mutation in the *FLO8* gene (*flo8-1*) that produces a truncated transcription factor altering expression of *FLO 11* (Liu *et al.* 1996). The initial results from our screens suggested a role for this truncated protein and were further confirmed by measuring the levels of mRNA in the *flo8*Δ strain (Figure-6). This deletion mutant displayed robust changes in all *TPK* and *BCY1* mRNAs, with the greatest effects on *TPK3* and *BCY1.* Our findings are in agreement with previous work reporting mRNA changes in the *flo8*Δ mutant for all TPKs and BCY1, analyzed using microarray technology (Hu *et al.* 2007). Interestingly, it has been recently reported that the C-terminal region of Flo8 has a novel transcriptional activation domain with a crucial role in activating transcription (Kim *et al.* 2014).

**Figure 6.**
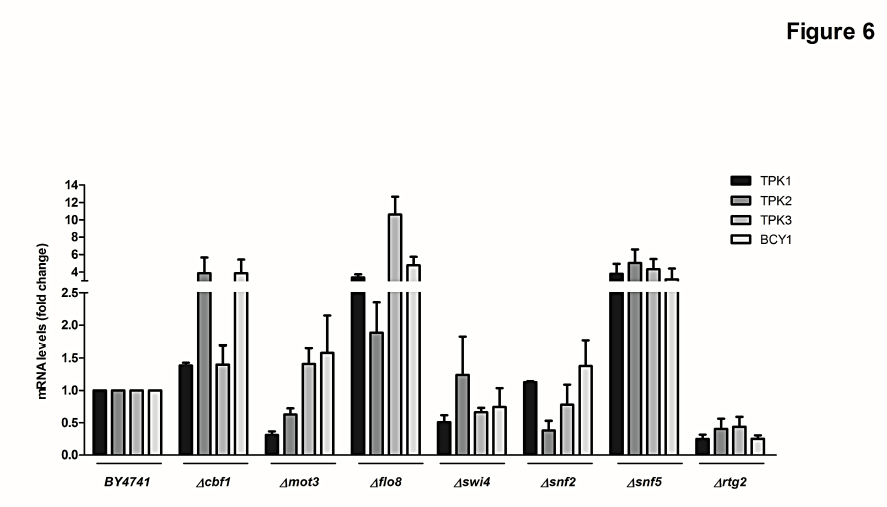
*TPKs* and *BCY1* endogenous mRNA levels were determined in WT (BY4741), *cbf1Δ, mot3Δ, flo8Δ, swi4Δ, snf2Δ, snf5Δ* and *rtg2Δ* strains by qRT-PCR and normalized to *TUB1* mRNA. The mRNA level for each subunit in the WT strain was defined as 1.

In addition other ategories related to mitochondria, vacuole, amino acids metabolism, transport and sporulation also affected the expression of all PKA subunits. While it is well documented for example that the cAMP pathway regulates mitochondria function as unregulated PKA activity can lead to the production of mitochondria that are prone to the production of ROS, and to an apoptotic form of cell death (Feliciello *et al.* 2005; Aun *et al.* 2013; Galello *et al.* 2014). Other processes as the progression of sporulation are known to be regulated from an interconnected signaling network that includes Ras/cAMP pathways altered even on rich media (Honigberg and Purnapatre 2003). Taking into account the reported antecedents mentioned above is clear how PKA activity regulates sporulation and mitochondria functioning, however it is not clear how the dysfunction of these processes would deregulate the expression of PKA subunits. Recently published results from our laboratory (Pautasso and Rossi 2014) demonstrated that the *TPK1*, *TPK2*, *TPK3* and *BCY1* promoter activities are upregulated during the stationary phase of growth in comparison with logarithmic phase. Additionally we have also observed an upregulation of the promoters of PKA subunits during growth in the presence of glycerol as carbon source, (results to be published elsewhere). Thus, during a respiratory metabolism, when a proper mitochondrial functioning is necessary, there is an activation of the PKA subunit promoters. Here, our SGA results are suggesting that defective mitochondria deregulate PKA subunits promoter activity. Therefore a reciprocal regulation could be proposed by which PKA regulation plays a physiological role in coordinating respiratory function with nutritional status in budding yeast and the respiratory metabolism would upregulate PKA subunit promoter activities.

Another interesting example is the regulation of vacuolar and proton acidification. Mutants lacking several subunits of the vacuolar-ATPase (*vma2*, *vma4*, *vma5*, *vma7*, *vma13*, *vma21* and *vma22*) have been identified in our screen. The V-ATPase is a proton pump that regulates cytosolic pH by pumping protons into the vacuole. Recent studies suggest that the V-ATPase is required for the glucose-mediated stimulation of PKA and that cytosolic pH serves as a second messenger in this regulatory pathway (Dechant and Peter 2010). Activation of V-ATPase is required for full activation of PKA upon glucose stimulation, thereby transducing, at least in part, the pH signal to PKA. As PKA activity autoregulates the activity of its own subunit promoters, malfunction of V-ATPase could indirectly impact the activity of the promoters by affecting the kinase activity.

## CONCLUSIONS

Regulation of eukaryotic gene expression is controlled at multiple levels, each distinguished by different spatial and temporal arrangements. The combination and orchestration between regulatory mechanisms at various levels are central to a precise gene expression pattern, which is essential to many critical biological processes. Kinases are sensitive to environmental perturbations and have different functions under different growth conditions. A kinase has many substrates and the signal specificity is key to achieve the exact and appropriate response to a given stimulus. In the cAMP-PKA pathway, the regulation of signal specificity is attained at several levels, one of them is the regulation of PKA subunits transcription. Our results showed an integrate network of common and singular regulators pathways of the transcription of each PKA subunit supporting the key role of the expression regulation of each subunit to achieve the final specificity in the cAMP-PKA signaling to sense and respond to environmental stimuli. Altogether, the results of our screen were consistent with previous findings and successful in identifying some of the expected modulators, while pointing to inositol and inositol-polyphosphates, choline and phosphate as novel upstream signals that regulate transcription of PKA subunit genes. Our results also open the possibility of a regulatory loop where pathways regulated by PKA phosphorylation control in turn, the level of expression of different PKA isoforms.

## Acknowledgments

We thank Dr Brenda Andrews (University of Toronto, Canada) for the generous gift of the BY4256 strain and BA1926 plasmid to perform the high-throughput functional assay. We also thank Dr Mauricio Terebiznik (University of Toronto at Scarborough, Canada) for access to the PharosFX Molecular Imager. This work was supported by grants from the Agencia Nacional de Promoción Científica y Tecnológica (PICT 2008-2195), from the University of Buenos Aires (UBA 2011-2014, 20020100100416), and from the Consejo Nacional de Investigaciones Científicas y Técnicas (CONICET) (PIP5239) to SR and by operating grants from the National Sciences and Engineering Research Council (NSERC) to GC and VZ. CP is a PhD fellow from CONICET (Argentina) and a scholar of the Emerging Leaders in the Americas Program (ELAP), supported by the Canadian Bureau for International Education.

---

**Supplemental Figure 1**- Distribution of log2 GFP:RFP ratios from genome-wide analysis of the *TPK1*, *TPK2 TPK3* and *BCY1* promoters grown in SD medium. The y-axis represents log2 GFP:RFP ratios measured from each deletion mutant displayed on the x-axis. The promoters-GFP fluorescence intensities are standardized to the control RPL39pr-RFP intensities.

**Supplemental Table 1**- Functional categorization of genes regulators identified in the screening for *TPK1*, *TPK2*, *TPK3* and *BCY1* promoters and of all genes in *S.cerevisiae.*
Classifications were performed based on the categories defined in the GO biological process, GO molecular function and MIPS databases using FunSpec.

